# iPSC-derived hepatocytes accurately recapitulate population diversity in alpha-1-antitrypsin deficiency and offer a novel *in vitro* model for large-scale drug efficacy screening studies

**DOI:** 10.1101/2025.01.21.634083

**Authors:** Carlos Gil, Vanesa Papastavrou, Gemma Gatti, Samuel Chung, George Kiloh, Ka Cheung, Magdalena Łukasiak, Chloe Robinson, Lia Panman, Ioannis Kasioulis, Nikolaos Nikolaou

## Abstract

**Background:** Alpha-1 antitrypsin deficiency (A1ATD) is a hereditary recessive disorder caused by mutations in the *SERPINA1* gene. It is a clinically under-recognised disease characterised by low circulating A1AT levels and intracellular accumulation of misfolded A1AT in hepatocytes. Deposition of excessive abnormal A1AT in the liver leads to liver failure, yet no specific treatments are available due to the lack of physiologically relevant disease modelling platforms.

**Methods:** We have hypothesised that human induced pluripotent stem cell (iPSC)-derived hepatocytes can provide an efficient platform to study A1ATD. Using CRISPR/Cas9, we have generated wild-type and A1ATD iPSC-derived hepatocytes (Opti-HEP) from healthy and A1ATD donors and developed a bioassay that mimics the accumulation of misfolded A1AT in the liver. Responses to the reference drug carbamazepine (CBZ), known to reduce intracellular misfolded A1AT levels, and RNA-based therapeutics were subsequently investigated.

**Results:** All lines successfully differentiated into hepatocytes as measured by comparable key hepatic and disease markers to those seen in primary human hepatocytes. The diseased lines displayed increased intracellular accumulation of misfolded A1AT compared to isogenic controls. Diseased cell lines showed significant decreases in intracellular accumulation of polymeric A1AT following transfection with RNA-based therapeutics, but a differential response upon treatment with CBZ.

**Conclusion:** We have developed a specific and robust *in vitro* model of A1ATD that recapitulates disease pathophysiology and responds to small molecule-based treatments and advanced therapeutic strategies. These data demonstrate the suitability of this model for large-scale efficacy screening studies for the treatment of A1ATD and help pave the way towards the development of novel therapies.

## 1. Introduction

Alpha-1 antitrypsin deficiency (A1ATD) is a clinically under-recognised genetic disorder that causes the defective production of alpha-1 antitrypsin protein (A1AT). A1AT, a 52 kDa protein encoded by the *SERPINA1* gene, is a liver-specific protease inhibitor responsible for the protection of host tissues by inhibiting neutrophil protease activity during inflammation [1]. Genetic mutations in the *SERPINA1* gene, also known as serpinopathies, cause A1ATD; amongst these, the most prevalent is the homozygous missense mutation p.E342K (or p.E366K), which is characterised by a single amino acid substitution where glutamic acid is replaced with lysine and is associated with >90% of all known clinical cases of severe A1ATD [2]. This mutation predisposes misfolding of the protein and promotes its polymerisation and accumulation in the endoplasmic reticulum of hepatocytes, causing a reduction in secretion of native protein into the plasma [3]. The disease manifests mainly in the lungs and liver, with chronic obstructive pulmonary disease being the most frequently reported condition. Clinical severity of liver disease in patients with A1ATD is extremely variable across patients and it ranges from fully asymptomatic to chronic hepatitis with or without cirrhosis [4]. Augmentation therapy is the most widely used therapy for patients with A1ATD, however, there are no current therapies targeting the hepatic manifestation of the disease other than liver transplant. Therefore, providing a physiologically relevant platform to identify novel therapeutics capable of alleviating the disease phenotype is of high interest.

The lack of an accurate model system that faithfully reproduces human hepatocyte biology has hindered the development of reliable platforms to screen potential therapeutics for A1ATD. Previous studies have focused either on genetically modified animal models of A1ATD or CHO cells over-expressing the polymer-forming version of the A1AT protein [5–7]; however, animal models fail to display the disease phenotype, whilst CHO cells are of non-human and liver origin, therefore lacking hepatic-enriched proteins that are required to ensure reliability during early drug screening and downstream clinical success. Primary hepatocytes from patients with A1ATD could offer a promising alternative, however, these are hard to source and culture, limiting their use for screening libraries of small molecules until the final stages of compound assessment. More recently, iPSC-derived hepatocytes have shown potential in bridging the gap and offer a suitable platform for the identification of new treatments against A1ATD. Indeed, a few studies have previously shown successful differentiation of patient-derived iPSCs harbouring mutations in the *A1AT* gene towards hepatocytes with increased accumulation of polymeric A1AT compared to wild-type controls, and reduction of polymer inclusions following TALEN-based gene editing or treatment with autophagy-inducing agents [8–10].

Despite the progress, there is still a critical need to complement these existing models and develop efficient platforms to identify novel compounds and therapeutics (e.g., gene therapeutic strategies) capable of mitigating A1ATD-associated pathology. Here, we extend the work on iPSC-derived hepatocytes and directly test the impact of previously used autophagy-inducing agents and novel RNA-based therapeutics on the A1ATD phenotype as well as the potential effect of genetic variation in drug efficacy using three distinct A1ATD patient-specific iPSC lines alongside a novel CRISPR/Cas9-derived A1ATD model.

## 2. Materials and methods

### 2.1. iPSC culture

Ethics for the wild-type (IPS-003) and patient 1 (IPS-001) DefiniGEN iPSC lines used in this work were approved under Addenbrooke’s Hospital reference 08/H0311/201. The A1ATD Patient 2 (#PiZZ1; IPS-039) and Patient 3 (#PiZZ6; IPS-041) iPSC lines as well as their corrected cell lines (# PiZZ-MM; #PiZZ6-MM) were received from Boston University; all material were generated with funding from the National Center for Advancing Translational Sciences (NCATS) at the NIH under grant number U01TR001810. iPSCs were cultured in Vitronectin XF-(#07180; STEMCELL Technologies, Cambridge, UK) coated tissue culture plastic using TeSR-E8 Medium (#05990; STEMCELL Technologies, Cambridge, UK) in a humidified tissue culture incubator at 37°C, 5% CO_2_. Cell medium was changed daily, and cells were passaged every 5-7 days using ReleSR (#100-0483; STEMCELL Technologies, Cambridge, UK).

### 2.2. iPSC differentiation and Opti-HEP culture

Direct differentiation of human iPSCs towards hepatocytes was performed as previously described with modifications [11]. To initiate differentiation, iPSCs were dissociated using StemPro Accutase Cell Dissociation Reagent (#A1110501; Life Technologies, Waltham, USA) and plated onto gelatine-coated dishes at a density of 20,000 cells/cm^2^. Following completion of differentiation, cells were dissociated using TrypLE (#12604013; Life Technologies, Waltham, USA) and replated onto freshly coated Collagen I Rat Protein (Gibco) 96-well plates. For monolayer Opti-HEP culture, cells were kept seeded at a density of 234,000 cells/cm^2^. For CBZ treatments (#C0270; Cambridge Bioscience, Cambridge, UK), drug was added in a range of concentrations (500 µM - 1.95 µM) 24 hours post-seeding, and cells were incubated for 72 ± 1 hours.

### 2.3. CRISPR/Cas9 gene editing

To generate the homozygous single nucleotide mutation E342K (E366K) in the *SERPINA1* gene, a 20-nucleotide PAM-NGG single-guide RNA (sgRNA) was designed to target exon 5. Repair template single stranded oligodeoxynucleotide (ssODN), containing the desired mutations, were designed with homologous genomic flanking sequences of 50 nucleotides length, centred around the targeted CRISPR/Cas9 cleavage site. Cas9/sgRNA (1/1.5 RNP complex) and 5 μM of single-stranded donor oligonucleotides (ssODN) were delivered to 1×10^6^ single cells by nucleofection using Lonza 4D nucleofection system (#V4XP-3024; Lonza Biologics, Slough, UK) according to the manufacturer’s protocol. Following nucleofection, cells were expanded and seeded in 96-well plates, before DNA was collected, extracted, and purified to identify positive clones via Sanger sequencing. The sequences for the sgRNAs, ssODNs, and genotyping primers are illustrated in Suppl. table 1.

### 2.4. Immunofluorescence

Cells were fixed with 4% p-formaldehyde (#PN28908; ThermoFisher Scientific, Waltham, MA) for 20 minutes, washed three times with DPBS (#D8537; Merck Life Science, Dorset, UK) and blocked for 1 h in 10% donkey serum (#S30-100ML; Merck Millipore, Dorset, UK) and 0.1% Triton X-100 (#X100-500ML; Merck Life Science, Dorset, UK) in DPBS. Following blocking, cells were incubated in DPBS containing 0.1% Triton X and 1% donkey serum with primary antibodies at 4°C overnight as follows: OCT4 (1:100; Santa Cruz Biotechnology Inc.; #SC5279), SOX2 (1:100; Merck/Millipore; #ab5603), NANOG (1:100; Cell Signalling; #D73G4), ALB (1:100; Abcam; #AB2406), total A1AT (1:100; Santa Cruz Biotechnology Inc.; #SC59438), HNF4A (1:200; Abcam; #AB41898), 2C1 A1AT(1:250; Hycult Biotech; #HM2289). Cells were washed 3 times with DPBS and incubated with respective secondary antibodies for 1 hour as follows: donkey anti-mouse 488 (1:1000; ThermoFisher Scientific; #A21202), donkey anti-rabbit 568 (1:1000; ThermoFisher Scientific; #A10042), DAPI (1:10000; Merck Life Science, Dorset, UK; #10236276001). After washing three times with DPBS for 5 minutes, cells were imaged using a Cellinsight CX7 High Content Analysis Platform (ThermoFisher Scientific, Waltham, MA).

### 2.5. Transfection and transduction studies

For siRNA transfection studies, cells were transfected with 100 nM ON-TARGETplus Human SERPINA1 (#5265, Horizon Discovery, Waterbeach, UK) siRNA – SMARTpool or ON-TARGETplus Non-targeting Pool (#D-001810-10, Horizon Discovery, Waterbeach, UK) as scrambled control. RNAimax was used at a concentration of 0.6 µl/well, and RNA samples were collected 48 hours post-transfection. For mRNA transfection studies, two concentrations (250 ng; 500 ng) of eGFP mRNA (#130-101-114, Miltenyi Biotec, Surrey, UK) were used using MessengerMAX (#LMRNA001; ThermoFisher Scientific, Waltham, MA) and GFP expression was investigated in a time-dependent manner (24-72 hours). For DNA transfection studies, cells were transfected with a GFP construct (pMax-GFP; 1.0 μg/μl; Lonza, Little Chesterford, UK) using DNA-Lipofectamine™ 3000 (#L3000001; ThermoFisher Scientific, Waltham, MA) at two different concentrations (100 ng; 500 ng), and GFP expression was investigated in a time-dependent manner (48-144 hours).

For AAV, transduction at two MOI (50,000 and 100,000) of AAV3-CAG-EGFP (#7108; Vector Biolabs, PA, USA) was performed, and GFP expression investigated in a time-dependent manner (3-12 days). For lentiviral particles, transduction at two MOIs (0.5, 1) of CMV-RFP-GFP-lentivirus (#CRUtest; Cellecta, CA, USA) was performed, and GFP expression was investigated in a time-dependent manner (3-12 days). Cells were stained with DAPI (1:10000; Merck Life Science, Dorset, UK; #10236276001) for nuclei quantification.

### 2.6. Gene expression analysis

Total RNA was extracted using ReliaPrep RNA Cell Miniprep System (#Z6012; Promega, Wisconsin, USA) according to the manufacturer’s protocol. Concentration was determined spectrophotometrically at OD_260_ on a Nanodrop spectrophotometer (ThermoFisher Scientific, Waltham, MA). Reverse transcription was performed using High-Capacity cDNA Reverse Transcription Kit (#4368813; Applied Biosystems, Waltham, MA).

All quantitative PCR experiments were conducted using an AB StepOne Plus sequence detection system. Reactions were performed in 10 μl volumes in 96-well plates in reaction buffer containing 5 μl TaqMan Fast Advanced Master Mix (#4444557; ThermoFisher Scientific, Waltham, MA). Life Technologies supplied all primers as predesigned Taqman Gene Expression Assays labelled with FAM and endogenous control (*PPIA*) with VIC. The relative expression ratio was calculated using the ΔΔCt method. Detailed information on the Taqman Gene Expression Assays can be found in Suppl. table 2.

### 2.7. Statistical analysis

Data are presented as mean ± standard error (SEM), unless stated otherwise. Statistical analysis was performed using paired or unpaired Student’s t-test. For comparison between different conditions, one-way analysis of variance (ANOVA) followed by Dunnett’s multiple comparison test was employed. Differences were regarded as significant if p-value was < 0.05. Statistical analysis was performed using GraphPad Prism (Graphpad Software Inc., La Jolla, USA).

## 3. Results

### 3.1. CRISPR/Cas9-derived A1ATD Opti-HEP demonstrate the disease phenotype *in vitro*

To model A1ATD *in vitro*, we initially performed iPSC quality control assessments to confirm suitability of our wild-type line for downstream differentiation towards Opti-HEP generation. Assessment of iPSC colony morphology and expression of pluripotency stem cell markers (OCT4, SOX2, NANOG) revealed compact and round iPSC colony shapes alongside >85% double positive cells expressing OCT4/NANOG and OCT4/SOX2 (Fig. 1A-D). Following confirmation of pluripotency, wild-type iPSCs were used to introduce the prevalent E342K mutation in exon 5 of the *SERPINA1* gene via CRISPR/Cas9 gene editing. Successful gene editing was confirmed by Sanger sequencing (Suppl. figure 1A) without any adverse effect on iPSC pluripotency or colony morphology (Fig. 1A-D). Both wild-type and A1ATD iPSC lines successfully differentiated in Opti-HEP as displayed by the characteristic cobblestone-like hepatocyte morphology (Fig. 1D), whilst mRNA expression analysis of the hepatocyte markers *ALB, A1AT*, and *HNF4A* revealed comparable mRNA levels between the two Opti-HEP lines and primary human hepatocytes (Fig. 1E). Following confirmation of hepatocyte function, we measured the protein levels of intracellular A1AT revealing a significant increase in both total and polymeric A1AT accumulation in A1ATD Opti-HEP compared to isogenic control (Fig. 1F, Suppl. figure 1B), confirming the presence of the disease phenotype in our model.

**Figure 1:**
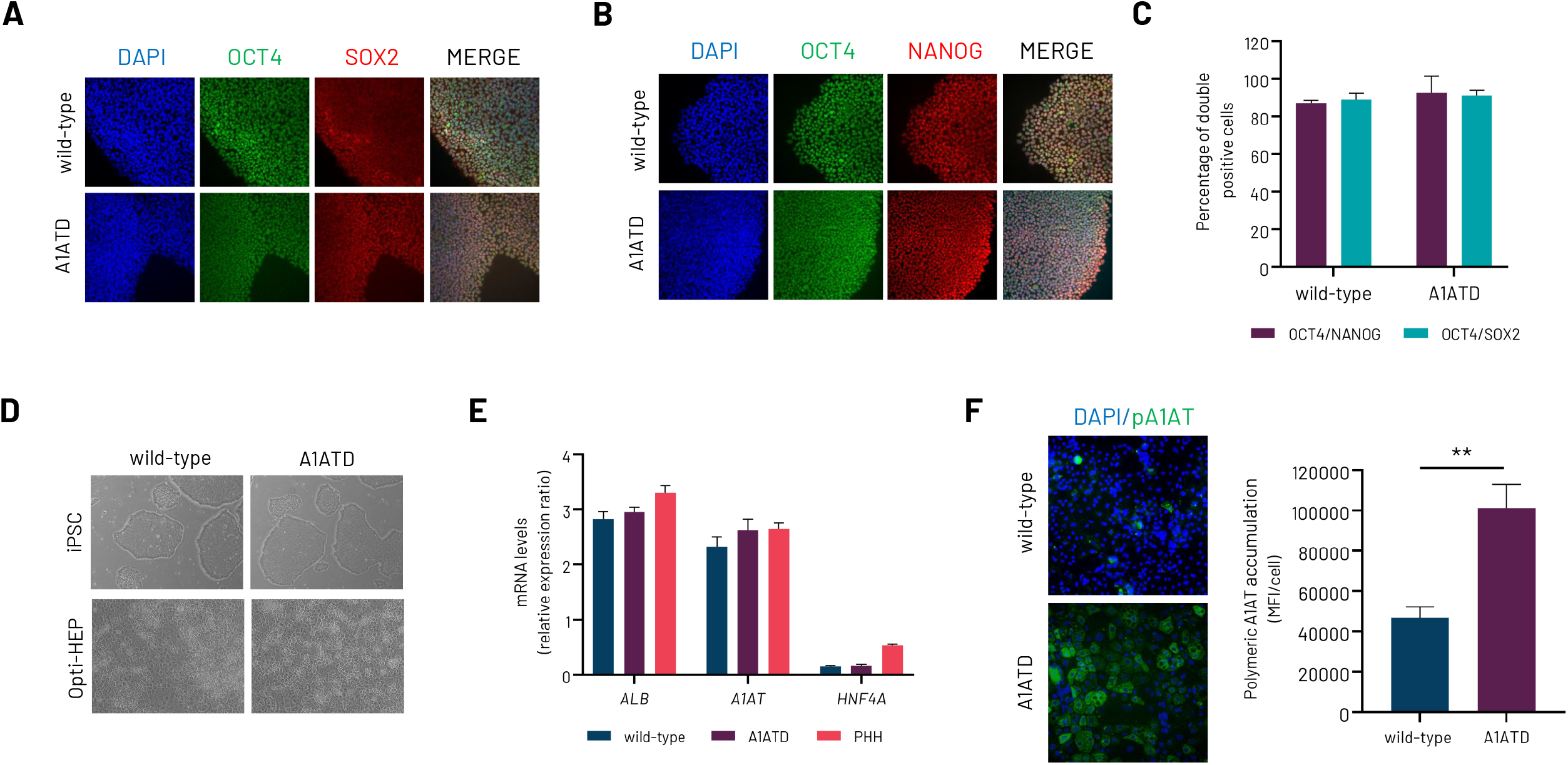
**(A-B)** Protein expression levels of Octamer-Binding Transcription Factor 4 (*OCT4*), SRY-Box 687 Transcription Factor 2 (*SOX2*), and Nanog Homeobox (*NANOG*) in wild-type induced pluripotent stem cells (iPSCs) (wild-type), and CRISPR/Cas9-derived alpha-1-antitrypsin iPSCs (A1ATD). **(C)** Percentage of double positive cells for OCT4/NANOG and OCT4/SOX2 in wild-type and A1ATD iPSCs. **(D)** Representative brightfield images of wild-type and A1ATD iPSCs and Opti-HEP demonstrating the round iPSC colony and hepatocyte cobblestone morphology, respectively. **(E)** mRNA expression levels of key hepatocyte markers albumin (*ALB*), alpha-1-antitrypsin (*A1AT*) and Hepatocyte Nuclear Factor 4 Alpha (*HNF4A*) in wild-type Opti-HEP, CRISPR/Cas9-derived A1ATD Opti-HEP, and primary human hepatocytes (PHH). **(F)** Immunostaining and quantification of intracellular levels of polymeric A1AT in A1ATD Opti-HEP compared to wild-type Opti-HEP. mRNA data were normalised to housekeeping gene Peptidylprolyl Isomerase A (*PPIA*). Nuclei were counterstained with DAPI (blue). Mean fluorescence intensity (MFI) was normalised to nuclei number. All results are presented as mean ± SEM of n=3 independent experiments. *p<0.05

### 3.2. Disease phenotype is reversed in A1ATD Opti-HEP using autophagy inducing agents and RNA-based therapeutics

Whilst there is currently no available treatment for A1ATD, previous studies have shown that the autophagy inducing agent CBZ can reduce intracellular A1AT aggregates both *in vitro* and *in vivo*, showing promise as a candidate therapeutic agent [12]. To this end, we tested efficacy of CBZ in A1ATD Opti-HEP by developing an endpoint assay where CBZ treatment was performed in a serial dilution ranging from 0-250 μM for 72 hours followed by measurement of intracellular polymeric A1AT accumulation using immunocytochemistry. In line with the data shown in figure 1F, the A1ATD phenotype was consistent and reproducible across the 96-well plate, with a coefficient of variation of 11.3% across three independent differentiation rounds (Suppl. figure 2A-B), and a dose-dependent decrease of polymeric A1AT was observed following CBZ treatment as shown in figure 2A.

**Figure 2:**
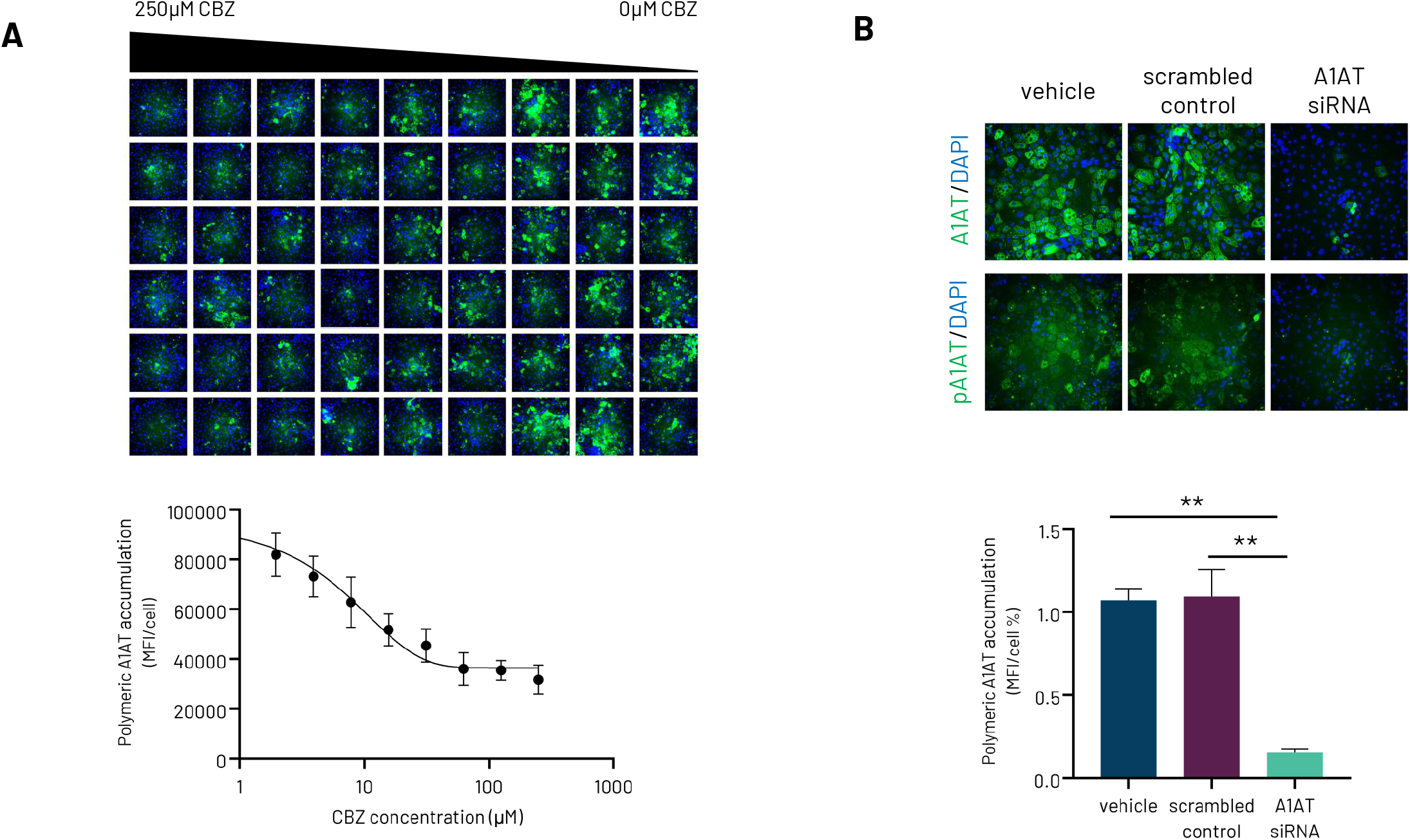
**(A)** Quantification and representative immunocytochemistry pictures of intracellular accumulation of polymeric A1AT in A1ATD Opti-HEP in response to treatment with different concentrations of carbamazepine (CBZ) following immunostaining with polymer-specific A1AT antibody. **(B)** Representative images of total and polymeric A1AT staining in CRISPR/Cas9-derived A1ATD Opti-HEP transfected for 72 hours with either vehicle, scrambled control, or siRNA targeting the *A1AT* gene. Nuclei were counterstained with DAPI (blue). Mean fluorescence intensity (MFI) was normalised to nuclei number. All results are presented as mean ± SEM of n=3 independent experiments. **p<0.01

RNA-based therapeutics have risen as an alternative and revolutionising method to treat diseases that cannot be targeted via the traditional drug development pipeline. To test whether our cells are additionally suitable for the screening of novel RNA-based therapeutics, A1ATD Opti-HEP were transfected with a short interference RNA (siRNA; 100 nM) targeting the *A1AT* gene for 72 hours. Immunocytochemistry analysis revealed a >70% decrease in both total and polymeric A1AT accumulation (Fig. 2B), confirming the suitability of our cells for the RNA-based therapeutic screening.

### 3.3. Patient-derived A1ATD Opti-HEP demonstrate the disease phenotype *in vitro* but differentially respond to autophagy inducing agents and RNA-based therapeutics

To investigate whether our differentiation platform is also applicable to patient-derived iPSCs harbouring the E342K mutation, a previously characterised A1ATD patient-derived iPSC line that has shown ability to demonstrate the disease phenotype *in vitro* was employed (Patient 1) [8,10]. Assessment of iPSC colony morphology and expression of pluripotency stem cell markers (OCT4, SOX2, NANOG) revealed compact and round iPSC colony shapes alongside >85% double positive cells expressing OCT4/NANOG, but low expression of SOX2 compared to our wild-type line (∼ 25% of either single or double positive cells expressing SOX2 or OCT4/SOX2, respectively) (Fig. 3A, Suppl. figure 3A). Additional analysis using a commercially available pluripotency assay service (Pluritest™, ThermoFisher Scientific) verified pluripotency of this iPSC line following screening against samples in the stem cell database, with a pluripotency score of 36.75 and novelty score of 1.33. Presence of the E342K mutation was confirmed by Sanger sequencing (Suppl. figure 3B). The Patient 1 iPSC line successfully differentiated to Opti-HEP, as shown by the characteristic cobblestone-like hepatocyte morphology and comparable mRNA expression levels of the mature hepatocyte markers *ALB, A1AT*, and *HNF4A* (Fig. 3B). Similar to our CRISPR/Cas9-derived A1ATD line, Patient 1 Opti-HEP successfully demonstrated the disease phenotype *in vitro*, as shown by significantly higher levels of intracellularly accumulated polymeric A1AT compared to our wild-type Opti-HEP (Fig. 3C). However, treatment of Patient 1 A1ATD Opti-HEP with increasing concentrations of CBZ had no effect in reducing polymeric A1AT levels (Fig. 3D). In contrast, and despite the lack of effect of the autophagy-inducing compound in rescuing the A1ATD phenotype, transfection of Patient 1 A1ATD Opti-HEP with the A1AT-specific siRNA (100 nM) resulted in a >80% decrease in both total and polymeric A1AT accumulation, as shown in figure 3E.

**Figure 3:**
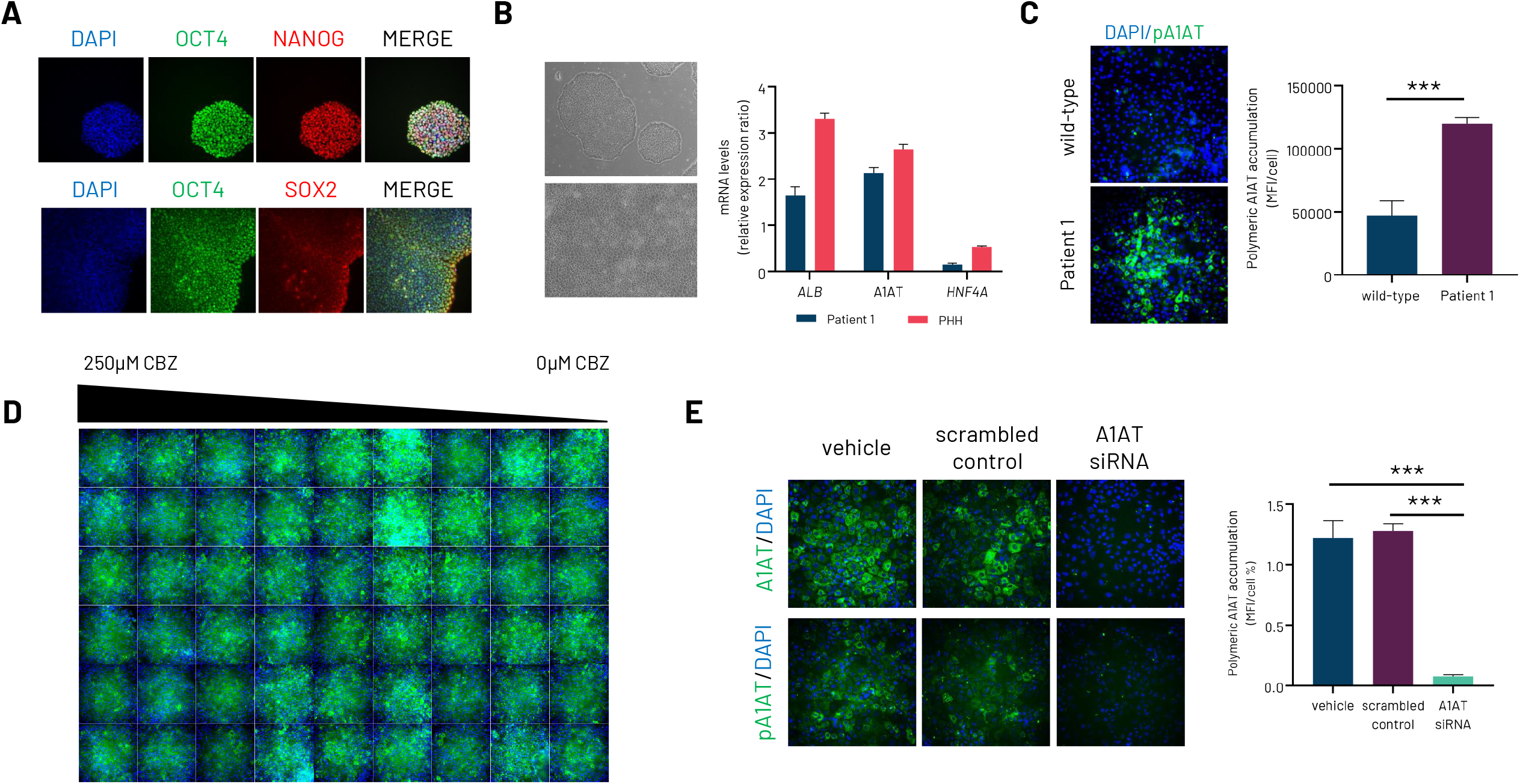
**(A)** Protein expression levels of Octamer-Binding Transcription Factor 4 (OCT4), SRY-Box 687 Transcription Factor 2 (SOX2), and Nanog Homeobox (NANOG) in patient 1-derived A1ATD iPSCs. **(B)** Representative brightfield images in Patient 1-derived A1ATD iPSCs and its respective Opti-HEP line demonstrating the round iPSC colony, hepatocyte cobblestone morphology, and mRNA expression levels of the key hepatocyte markers albumin (*ALB*), alpha-1-antitrypsin (*A1AT*) and Hepatocyte Nuclear Factor 4 Alpha (*HNF4A*) compared to primary human hepatocytes (PHH). **(C)** Immunostaining and quantification of intracellular levels of polymeric A1AT in Patient 1-A1ATD Opti-HEP compared to wild-type Opti-HEP. **(D)** Intracellular accumulation of polymeric A1AT in Patient 1-A1ATD Opti-HEP in response to treatment with increasing concentrations of carbamazepine (CBZ) following immunostaining with polymer-specific A1AT antibody. **(E)** Representative pictures of total and polymeric A1AT staining in Patient 1-derived A1ATD Opti-HEP transfected for 72 hours with either vehicle, scrambled control, or siRNA targeting the *A1AT* gene. Nuclei were counterstained with DAPI (blue). Mean fluorescence intensity (MFI) was normalised to nuclei number. All results are presented as mean ± SEM of n=2-3 independent experiments. ***p<0.001

### 3.4. Additional patient-derived A1ATD Opti-HEP demonstrate the disease phenotype *in vitro* and successfully respond to autophagy inducing agents

Informed by the data presented in figure 3, we wanted to investigate whether the observed differences between our CRISPR/Cas9- and Patient 1-derived Opti-HEP in response to CBZ are driven by genetic background variation or technical limitations in our platform. To achieve this, two additional homozygous patient-derived iPSC lines (Patient 2, Patient 3) were employed alongside their CRISPR-edited corrected controls. Presence of the E342K mutation as well as successful correction of the mutation was confirmed by Sanger sequencing (Suppl. figure 4A-B). Similar to our wild-type and Patient 1 iPSC lines, assessment of iPSC colony morphology and expression of OCT4, SOX2, and NANOG revealed compact and round iPSC colony shapes alongside >85% double positive cells expressing OCT4/NANOG and OCT4/SOX2 in all patient-derived iPSC lines (Fig. 4A-D). All iPSC lines successfully differentiated into Opti-HEP as displayed by the characteristic cobblestone-like hepatocyte morphology (Fig. 4D). Finally, measurement of both total and polymeric A1AT accumulation showed significantly higher levels in the two patient-derived Opti-HEP lines compared to their isogenic controls consistent with the disease phenotype (Fig. 4E; Suppl. figure 4C-D).

**Figure 4:**
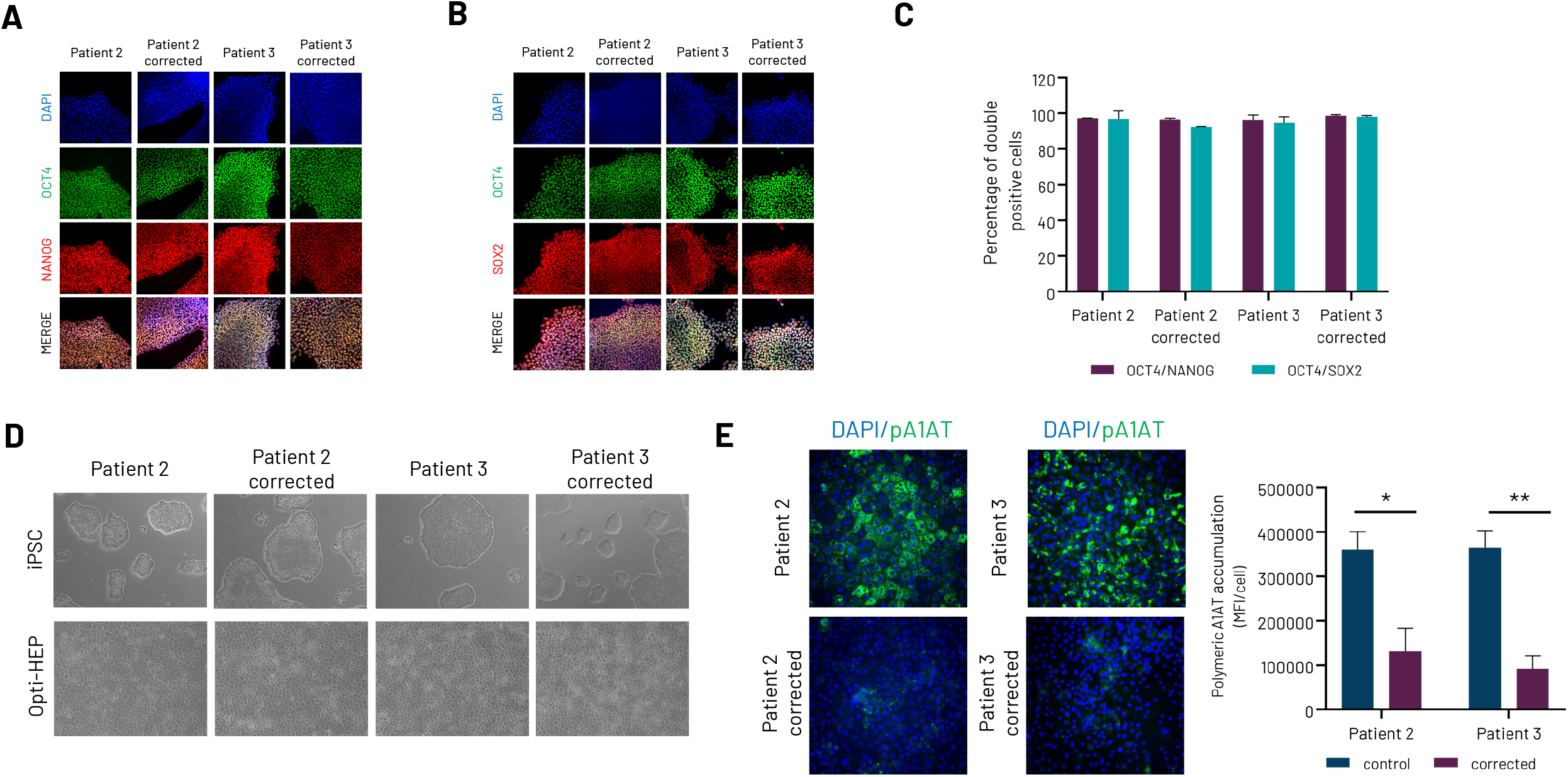
**(A-B)** Protein expression levels of Octamer-Binding Transcription Factor 4 (*OCT4*), SRY-Box 687 Transcription Factor 2 (*SOX2*), and Nanog Homeobox (*NANOG*) in patient-derived A1ATD iPSCs as well as their CRISPR/Cas9-corrected isogenic controls. **(C)** Percentage of double positive cells for OCT4/NANOG and OCT4/SOX2 in patient-derived A1ATD iPSCs and CRISPR/Cas9-corrected isogenic controls. **(D)** Representative brightfield images in patient-derived A1ATD iPSCs, CRISPR/Cas9-corrected isogenic controls and their respective Opti-HEP lines demonstrating the round iPSC colony and hepatocyte cobblestone morphology. **(E)** Immunostaining and quantification of intracellular levels of polymeric A1AT in patient-derived A1ATD Opti-HEP compared to their CRISPR/Cas9-corrected isogenic controls. Nuclei were counterstained with DAPI (blue). Mean fluorescence intensity (MFI) was normalised to nuclei number. All results are presented as mean ± SEM of n=3 independent experiments. *p<0.05, **p<0.01

Same as above, response of patient-derived A1ATD Opti-HEP to potential therapeutic compounds was also evaluated following 72-hour treatment with CBZ in a dose-dependent manner. Prior to any treatment, reproducibility and batch-to-batch variation were examined with the cells demonstrating a uniform A1ATD phenotype across the 96-well plate and a coefficient of variation of 10.1% and 12% for Patient 1 and Patient 2 Opti-HEP, respectively (Suppl. figure 5A-B). Crucially, both patient Opti-HEP lines showed a consistent, dose-dependent response to CBZ, as measured by decreasing intracellular levels of polymeric A1AT across increasing drug concentrations (Fig. 5A-B), highlighting the additional ability of our model to address donor-to-donor variability in drug efficacy screening.

**Figure 5:**
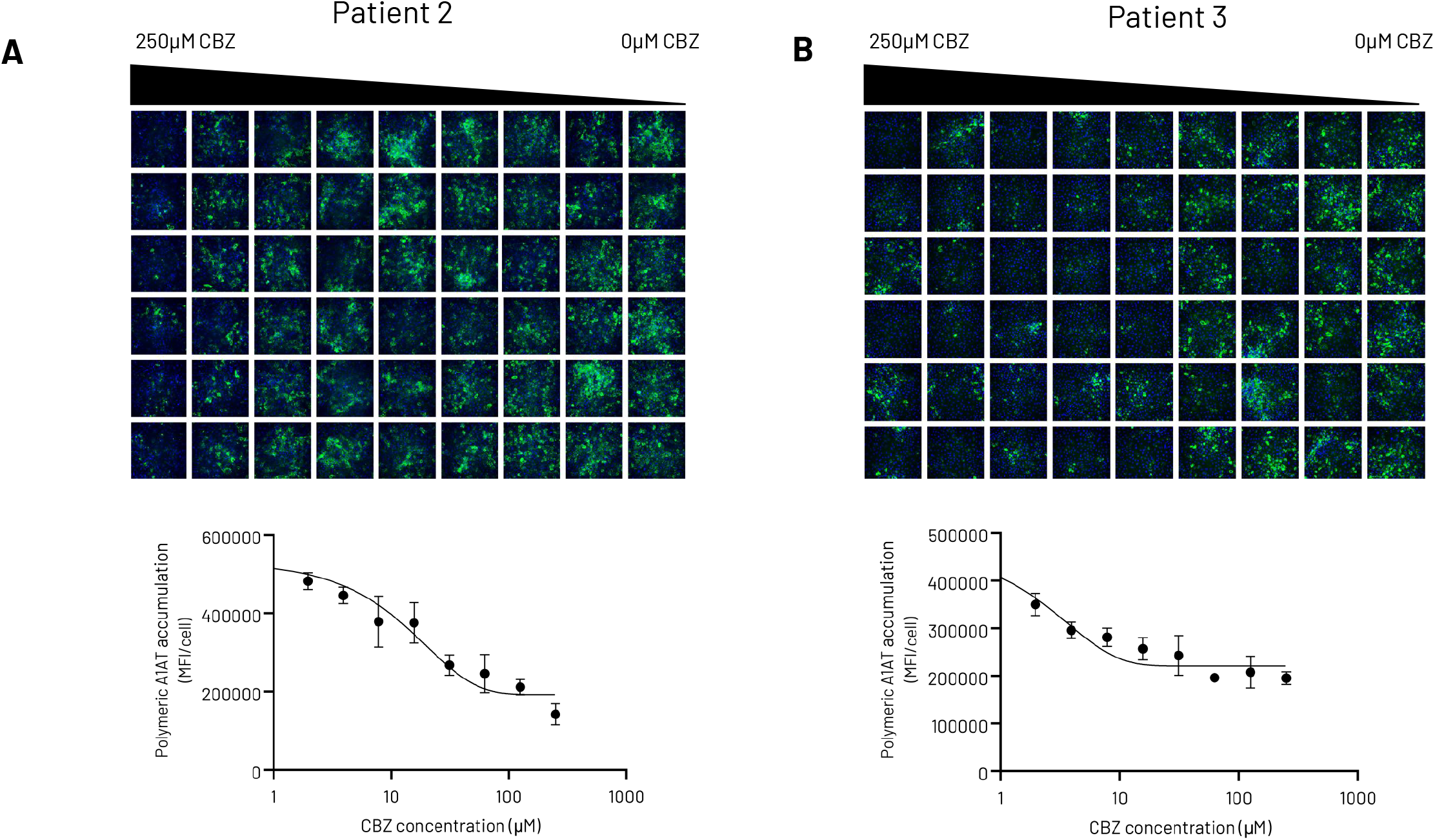
**(A)** Quantification and representative immunocytochemistry pictures of intracellular accumulation of polymeric A1AT in Patient 2 A1ATD Opti-HEP in response to treatment with increasing concentrations of carbamazepine (CBZ) following immunostaining with polymer-specific A1AT antibody. **(B)** Quantification and representative immunocytochemistry pictures of intracellular accumulation of polymeric A1AT in Patient 3 A1ATD Opti-HEP in response to treatment with increasing concentrations of carbamazepine (CBZ) following immunostaining with polymer-specific A1AT antibody. Results expressed as mean ± SEM of n=4 independent experiments.

## 4. Discussion

In this study, we have combined the opportunities iPSC-derived hepatocytes and CRISPR/Cas9 offer in disease model development and demonstrated the generation of a novel *in vitro* platform for the large-scale screening of small compounds and RNA-based therapeutics for the treatment of A1ATD. Importantly, our work highlights the crucial effect of genetic variation in drug efficacy.

To date, at least 100 different variants in the *A1AT* gene have been described. Amongst these, the Glu342Lys mutation (E342K, also known as the Z variant) is the most prominent and results to a range of mild (in heterozygous patients – MZ) to strong (in homozygous patients – ZZ) clinical features, including decreased serum concentrations of A1AT and predisposition to both liver and lung diseases [13]. Additional variants, including the Glu264Val (S variant), Arg233Cys (F variant) and Arg39Cys (I variant) have been described and also related to a spectrum of clinical features, ranging from no clear phenotype to moderate predisposition to liver disease and lung emphysema [14]. In our study, we have focused on the E342K mutation; however, the applicability of our iPSC differentiation protocol to a variety of different clones highlights the potential for additional mutations to be studied through the generation of either CRISPR/Cas9-derived cell models or reprogramming of somatic cells from patients harbouring these variants. Indeed, a recent study by Kaserman et al. [15] have described the generation of heterozygous for the Z variant iPSC-derived hepatocytes, demonstrating a milder disease phenotype compared to the homozygous model.

The expandable nature of iPSCs and the physiological relevance of differentiated iPSC-derived hepatocytes dovetail to produce disease models ideally suited for therapeutic screening assays. Indeed, a variety of monogenic and multi-factorial diseases have been previously modelled using iPSC-derived hepatocytes [16,17], however, developing platforms that can either directly or indirectly measure disease phenotypes in a high-throughput format has been challenging. Batch-to-batch variability in addition to incomplete cell maturation and low-drug metabolising capacity have been previously reported as issues that hinder the use of iPSC-derived hepatocytes in large-scale drug efficacy and toxicity screening [16,18]. Addressing these limitations, we have recently reported the development of optimised iPSC-derived hepatocytes with improved metabolic and drug metabolising functions as well as the generation of liver disease models for rare monogenic diseases, including urea cycle disorders and familial cholestasis [19]; expanding this work, this study highlights not only the applicability of our platform in additional disease models, but also the high reproducibility of our assays, as evident by the low CoV values shown above.

Using CBZ as an exemplar compound against A1ATD, we have shown reducing intercellular polymer accumulation in a dose-dependent manner. Our data are consistent with previous studies showing the effect of the autophagy-enhancing agent in decreasing hepatic A1AT load and fibrosis in cell and rodent models [9,12]. Additional autophagy-enhancing agents and regulators have been screened with promising results [9,20,21], but the exact mechanism through which autophagy mediates the A1ATD phenotype requires further investigation [22]. More recently, a small molecule that inhibits the polymerisation of the Z A1AT protein and increases secretion *in vitro* was reported [6].

Emerging gene delivery therapeutic strategies, such as siRNA-based approaches, have been particularly attractive over the last years. These therapeutics offer a promising alternative for the treatment of diseases that cannot be cured with conventional drug groups, since they can target almost every genetic component within a cell, whilst their transient nature reduces the likelihood of permanent mutations in the patients’ genome [23,24]. Indeed, to date, at least three siRNA drugs have been approved by the FDA targeting a variety of diseases, including hereditary transthyretin amyloidosis, hepatic porphyria, and primary hyperoxaluria type 1 [25–27], whilst several novel siRNA candidates are currently under Phase III clinical trials [23]. Our work, where A1AT-specific siRNA was used to reduce polymeric intracellular A1AT levels shows that this class of novel therapeutics can be successfully validated using our platform, and this can be expanded to additional technologies, such as base editing and gene therapies. For A1ATD, two RNA editing candidates are currently under clinical development ([Wave Life Sciences Press Release](https://ir.wavelifesciences.com/news-releases/news-release-details/wave-life-sciences-announces-initiation-dosing-restoraation), [Korro Bio Press Release](https://ir.korrobio.com/news-releases/news-release-details/korro-announces-selection-its-first-development-candidate)). Of note, many of these newly made RNAi-based drugs are conjugated with N-acetylgalactosamine (GalNAc), a high-affinity ligand for the hepatocyte-specific asialoglycoprotein receptor (ASGR1), thereby ensuring successful and organ-specific delivery of the siRNA into the liver. In line with this, we have previously shown expression and functionality of ASGR1 in Opti-HEP using GalNAc-siRNA conjugates [19]; extending this data set, we are now showing that our cells can offer a suitable platform for additional gene delivery strategies, including adeno-associated viruses, lentiviruses, mRNA, or DNA plasmid (Suppl. figure 6A-D).

Our platform does not come without limitations. Despite the reproducible intracellular accumulation of polymeric A1AT in our diseased Opti-HEP and the subsequent response to either CBZ or A1AT-siRNA, no significant differences in A1AT secretion were observed (data not shown). It may be possible that the lack of an accurate ELISA-based assay able to distinguish total from polymeric A1AT levels is responsible for this issue; indeed, in-house efforts to develop an immunofluorescence-based assay able to specifically measure polymeric A1AT levels in the culture media of A1ATD cells pre- and post-treatment is currently ongoing. In addition, the development of additional *in vitro* assays and co-culture systems, including the use of neutrophils for the generation/inhibition of neutrophil elastase activity, would enhance our understanding on the crosstalk between different cell types for the identification of novel therapeutics against A1ATD.

To conclude, we have generated an efficient and reproducible *in vitro* model for the large-scale efficacy screening of novel therapies against A1ATD. Importantly, this platform offers the opportunity to address population diversity and detect potential differences in the efficacy of these therapeutics that are driven by genetic background early in the drug developmental pipelines. It is anticipated that by using models with enhanced predictivity at earlier stages of development, resources can be deployed with greater efficiency and reduce the number of treatments suffering damaging late-stage clinical trial failures and post-approval market withdrawals.

## List of abbreviations

AAV: adeno-associated virus
ALB: albumin
A1AT: alpha-1 antitrypsin
A1ATD: alpha-1 antitrypsin deficiency
CBZ: carbamazepine
DMSO: Dimethyl sulfoxide
GAPDH: Glyceraldehyde-3-Phosphate Dehydrogenase
HNF4A: hepatocyte nuclear factor 4A
iPSCs: induced pluripotent stem cells
MOI: multiplicity of infection
NANOG: Nanog Homeobox
OCT4: Octamer-Binding Transcription Factor 4
PHH: primary human hepatocytes
PPIA: Peptidylprolyl Isomerase A
PVDF: polyvinylidene difluoride
sgRNA: single-guide RNA
SOX2: SRY-Box Transcription Factor 2
ssODN: single stranded oligodeoxynucleotide

## Supplementary figure legends

**Suppl. figure 1: (A)** Sanger sequencing chromatogram showing wild-type and mutated induced pluripotent stem cells (iPSC) carrying the E342K mutation (GAG>AAG) in the *A1AT* gene. The codon change is highlighted with grey. **(B)** Representative images of immunostaining of total A1AT in CRISPR/Cas9-derived A1ATD Opti-HEP compared to isogenic control. Objective 10X. Nuclei were counterstained with DAPI.

**Suppl. figure 2: (A)** Representative immunocytochemistry images and quantification of polymeric A1AT in CRISPR/Cas9-derived A1ATD Opti-HEP across the central 60 wells of a 96-well plate relative to the average of the plate set as 1. **(B)** Quantification of intracellular levels of polymeric A1AT in CRISPR/Cas9-derived A1ATD Opti-HEP across the 60 central wells of a 96-well plate. Results expressed as average from n=3 independent experiments.

**Suppl. figure 3: (A)** Quantification of percentage of single and double positive Patient 1 iPSCs expressing Octamer-Binding Transcription Factor 4 (OCT4), SRY-Box 687 Transcription Factor 2 (SOX2), and Nanog Homeobox (NANOG). **(B)** Sanger sequencing chromatogram showing presence of the E342K mutation in Patient 1-derived A1ATD iPSCs compared to wild-type control. The codon change is highlighted with grey. Results are presented as mean ± SD of n=2 independent experiments.

**Suppl. figure 4: (A-B)** Sanger sequencing chromatograms showing A1ATD patient-derived (Patient 2 and 3) and corrected for the E342K mutation iPSCs. The codon change is highlighted with grey. **(C-D)** Representative images of immunostaining of total A1AT in patient-derived A1ATD Opti-HEP and their corrected isogenic controls. Nuclei were counterstained with DAPI. Objective 10X.

**Suppl. figure 5: (A-B)** Representative immunocytochemistry images and quantification of polymeric A1AT in patient-derived A1ATD Opti-HEP across the central 60 wells of a 96-well plate relative to the average of the plate set as 1.

**Suppl. figure 6:** GFP expression and quantification of GFP-positive cells in DefiniGEN Opti-HEP following transduction or transfection with **(A)** GFP-AAV (Serotype 3), **(B)** GFP-lentivirus, **(C)** GFP-mRNA and **(D)** GFP-DNA in a time-dependent manner. Results are expressed as mean ± SEM of n=3 independent experiments. Nuclei were counterstained with DAPI. Scale bar: 400 μm.

