## Supplementary figures for "iPSC-derived hepatocytes accurately recapitulate population diversity in alpha-1-antitrypsin deficiency and offer a novel *in vitro* model for large-scale drug efficacy screening studies"

### Suppl. figure 1

**A**

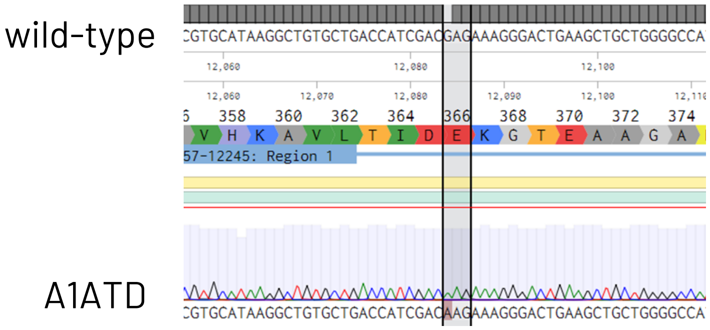


**B**

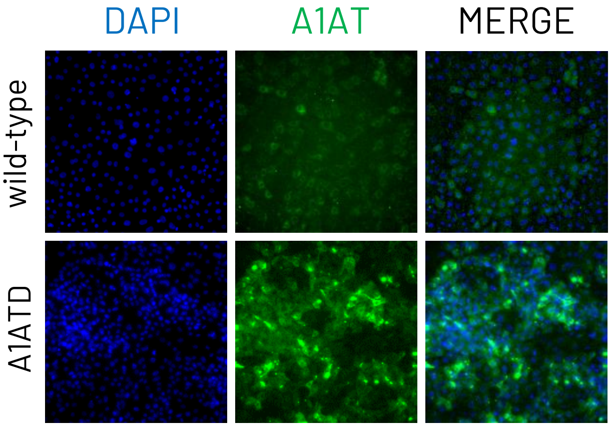

### Suppl. figure 2

**A**

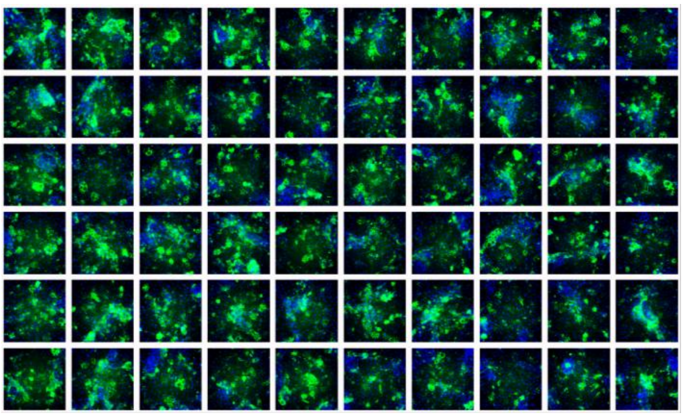

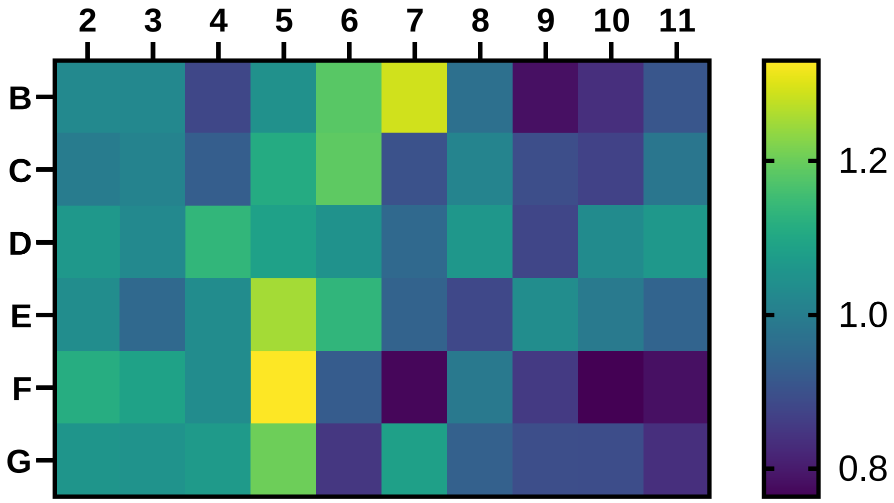


**B**

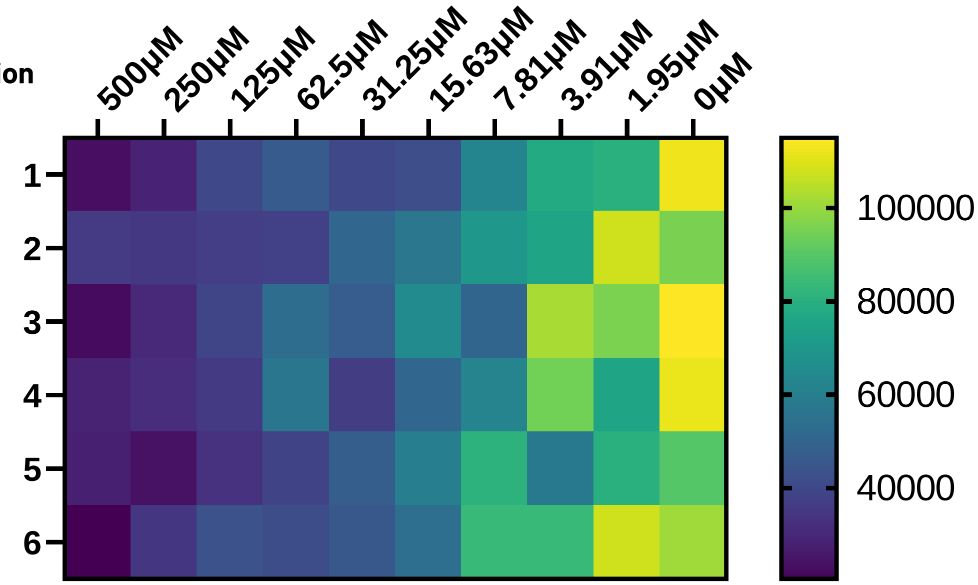

### Suppl. figure 3

**A**

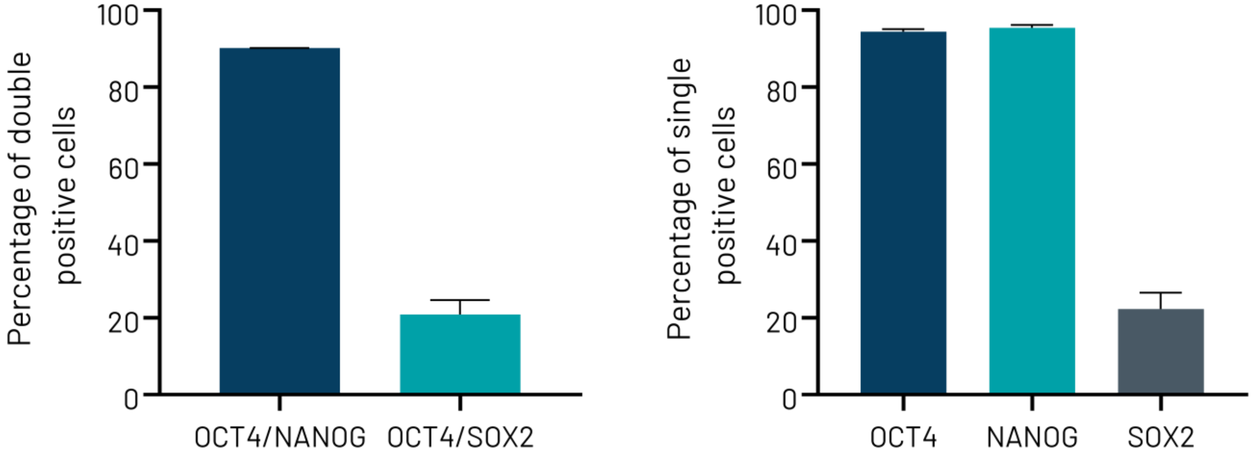


**B**

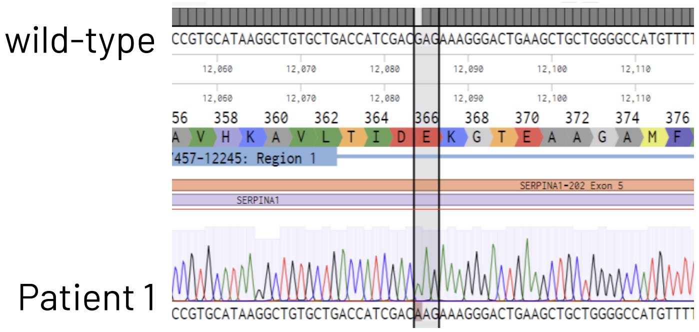

### Suppl. figure 4

**A**

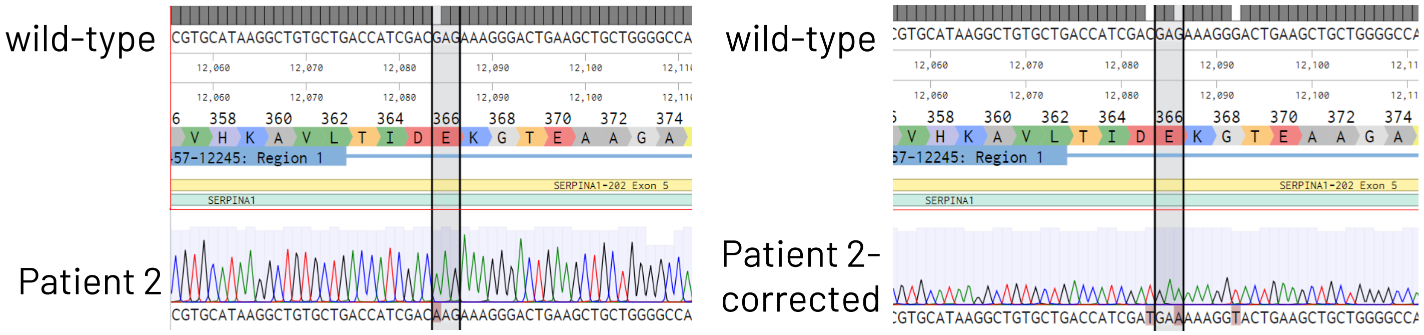


**B**

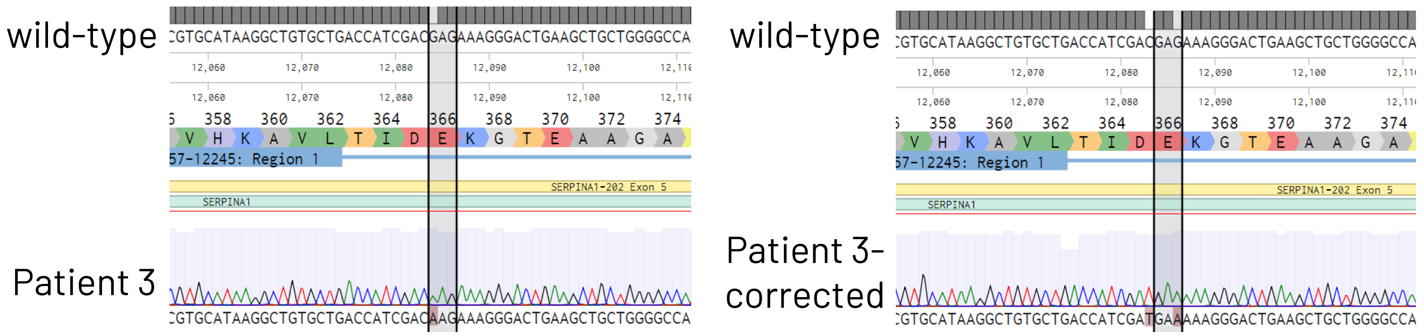


**C**

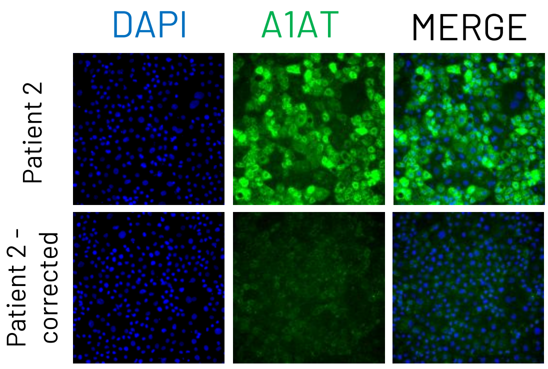


**D**

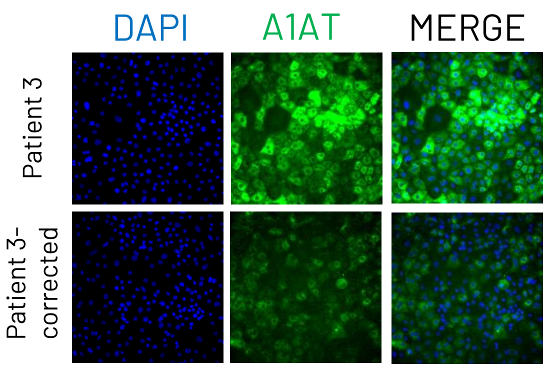

### Suppl. figure 5

A

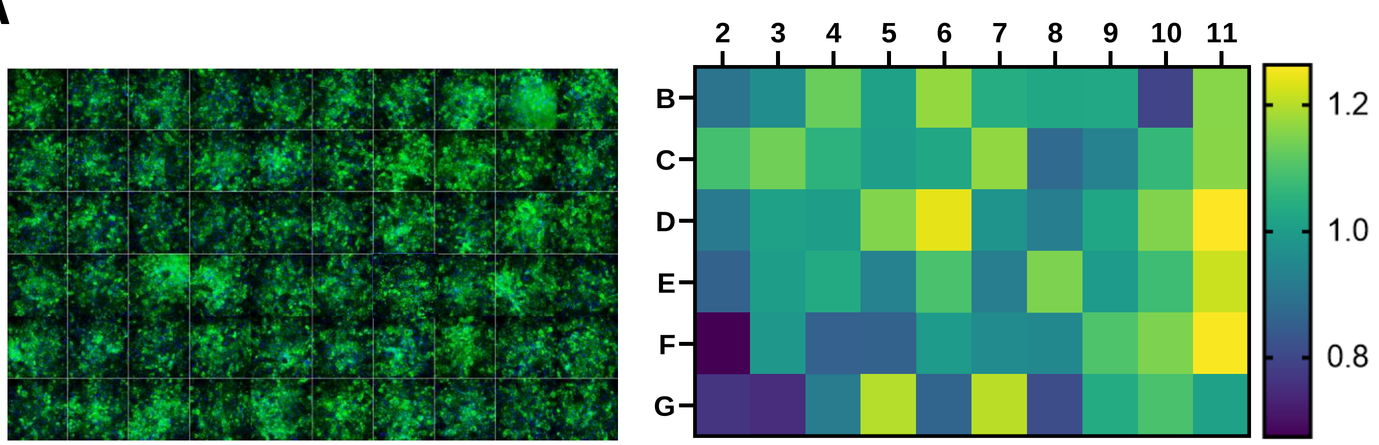


B

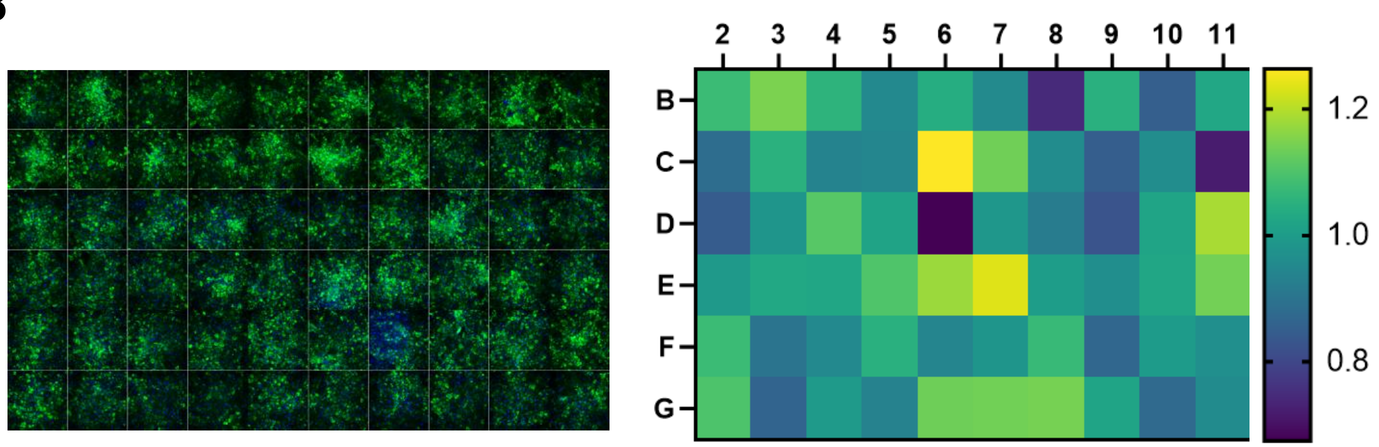

### Suppl. figure 6

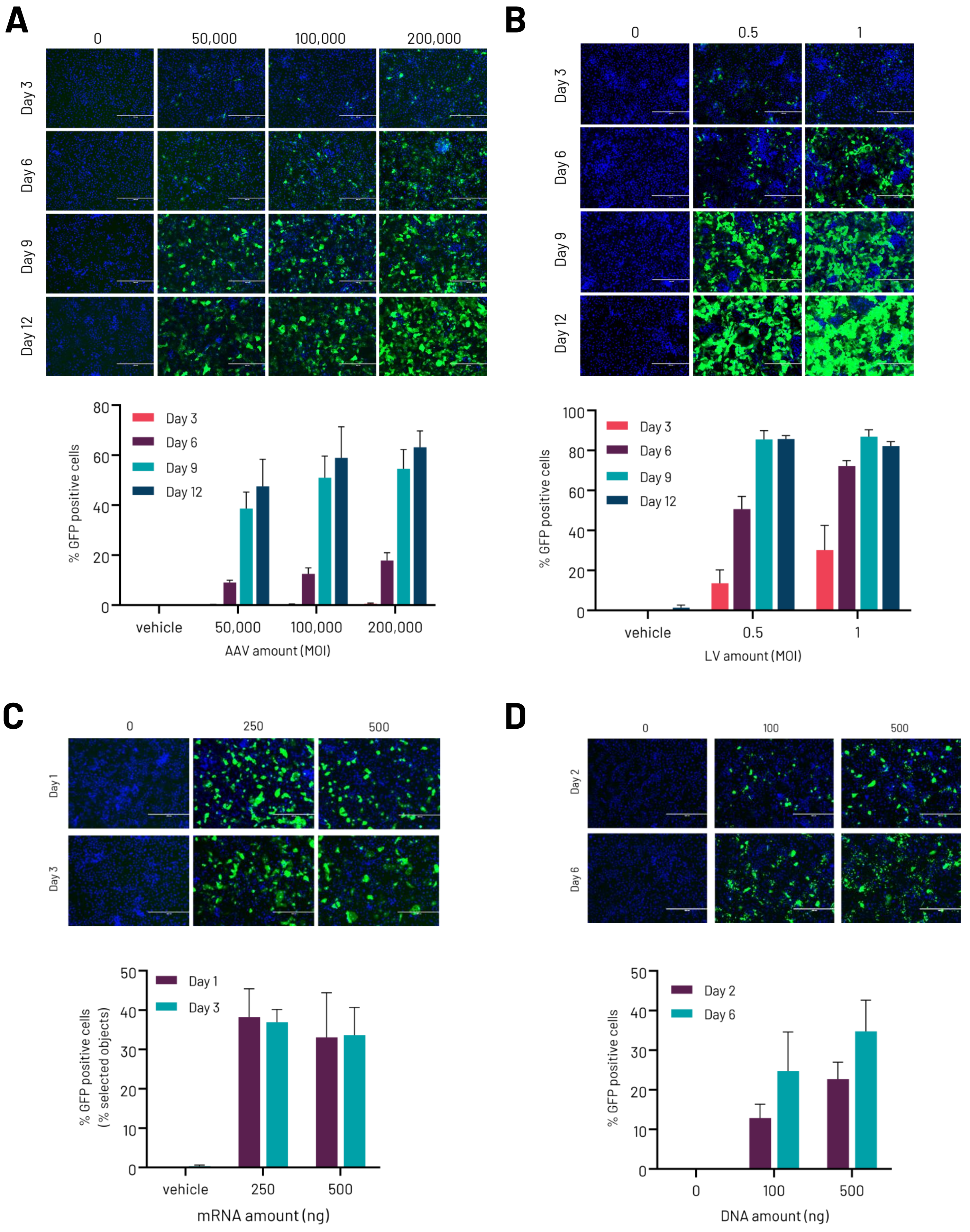

### Suppl. table 1

List of sgRNAs, ssODNs and genotyping primers used for CRISPR-Cas9 gene editing

| Target gene | Mutation | sgRNA Sequence | ssODN Sequence | Forward primer | Reverse primer |
|---|---|---|---|---|---|
| A1AT | E342K (Exon 7) | 5′-TGCTGACCATCGACGAGAAA-3′ | 5′-TCTGCTTCTCTCCCCTCCAGGCCGTGCATAAGGCTGTGCTGACCATCGAC**A**AGAAAGG**A**ACTGAAGCTGCTGGGGCCATGTTTTTAGAGGCCATACCCAT-3′ | 5′-GTCTGGGCACTGTGAAGGTC-3′ | 5′-GAGGGGTTGAGGAGCGAGAG-3′ |

### Suppl. table 2

List of TaqMan probes used for qPCR

| Target gene | Cat.No. | Dye | Probe ID |
|---|---|---|---|
| *PPIA* | 4326316E | VIC-MGB | Hs99999904_m1 |
| *ALB* | 4331182 | FAM-MGB | Hs00910225_m1 |
| *A1AT* | 4331182 | FAM-MGB | Hs01097800_m1 |
| *HNF4A* | 4331182 | FAM-MGB | Hs00230853_m1 |
